# Fixed single-cell RNA sequencing for understanding virus infection and host response

**DOI:** 10.1101/2020.09.17.302232

**Authors:** Hoang Van Phan, Michiel van Gent, Nir Drayman, Anindita Basu, Michaela U. Gack, Savaş Tay

## Abstract

Single-cell transcriptomic studies that require intracellular protein staining, rare cell sorting, or inactivation of infectious pathogens are severely limited because current high-throughput RNA sequencing methods are incompatible with paraformaldehyde treatment, a common tissue and cell fixation and preservation technique. Here we present FD-seq, a high-throughput method for droplet-based RNA sequencing of paraformaldehyde-fixed, stained and sorted single-cells. We show that FD-seq preserves the mRNA integrity and relative abundances during fixation and subsequent cell retrieval. Furthermore, FD-seq detects a higher number of genes and transcripts than methanol fixation. We applied FD-seq to investigate two important questions in Virology. First, by analyzing a rare population of cells supporting lytic reactivation of the human tumor virus KSHV, we identified *TMEM119* as a host factor that mediates viral reactivation. Second, we found that upon infection with the betacoronavirus OC43, which causes the common cold and is a close relative of SARS-CoV-2, pro-inflammatory pathways are primarily upregulated in lowly-infected cells that are exposed to the virus but fail to express high levels of viral genes. FD-seq thus enables integrating phenotypic with transcriptomic information in rare cell populations, and preserving and inactivating pathogenic samples that cannot be handled under regular biosafety measures.

## INTRODUCTION

Single-cell RNA sequencing (scRNA-seq) has found important biological applications, from discovery of new cell types^1^ to mapping the transcriptional landscape of human embryonic stem cells^2^. Droplet-based scRNA-seq technologies, such as Drop-seq^3^ and 10X Chromium^4^, are particularly powerful due to their high throughput: thousands of single cells can be analyzed in a single experiment. However, even with these high-throughput techniques, analyzing rare cell populations remains a challenging task, often requiring protein-based enrichment for the cell population of interest before scRNA-seq^5,6^.

Many cell types require intracellular protein staining to be enriched. For example, Foxp3 is an intracellular marker of regulatory T cells^7^, and Oct4 and Nanog are intracellular reprogramming markers of induced pluripotent stem cells^8^. Intracellular protein staining requires cell fixation, which is most commonly achieved with paraformaldehyde (PFA) or methanol fixation. Drop-seq and 10X Chromium have been shown to be compatible with methanol-fixed cells^9,10^, but not with PFA fixation. In many applications, PFA fixation is preferred over methanol fixation due to its better signal-to-noise ratio for intracellular staining^11,12^, and the improved preservation of fluorescent protein activity. scRNA-seq of PFA-fixed cells has only been achieved with a low throughput plate-based method^5^, severely limiting the applicability of this method to a wide range of problems that search for rare phenotypes in broad cellular populations. A high-throughput scRNA-seq method of PFA-fixed cells would enable the application of single cell analysis for many problems in signaling, immunity, development, stem cells, and infectious diseases.

Here we describe FD-seq (Fixed Droplet RNA sequencing), a droplet-based high-throughput RNA sequencing of PFA-fixed, stained and sorted single cells. We show that FD-seq preserves the RNA integrity and relative transcripts abundances compared to standard Drop-seq for live cells. We also show that FD-seq is superior to the methanol fixation protocol, yielding a higher number of detected genes and transcripts.

As a proof-of-concept, we applied FD-seq to study two important problems in Virology. First, we studied the low-level reactivation of Kaposi’s sarcoma-associated herpesvirus (KSHV) in tumor cells. KSHV, also known as human herpesvirus type 8 (HHV-8), is a human gammaherpesvirus that causes a number of malignancies such as Kaposi’s sarcoma, primary effusion lymphoma and multicentric Castleman’s disease^13,14^. There is a considerable interest in unraveling the molecular details of the host factors that modulate KSHV latency and reactivation, because both latency and low-level reactivation are known to contribute to viral tumorigenesis^15^, and therapeutic induction of reactivation could sensitize latently-infected cells to currently available anti-herpesvirus drugs^16^. Detailed analysis of KSHV reactivation, however, is currently limited by this restricted reactivation: only a small proportion of latently-infected cells typically undergoes reactivation, even when treated with known chemical inducing agents such as sodium butyrate (NaBut) and tetradecanoyl phorbol acetate (TPA)^13^. We hypothesized that the differences in the abundance of specific host factors between individual cells contribute to the propensity of latently KSHV-infected cells to enter lytic reactivation. Using FD-seq, we present the first single-cell transcriptomic analysis of human primary effusion lymphoma (PEL) cells undergoing reactivation. We found that in reactivated cells, the expression levels of viral genes were extremely heterogeneous, with some cells expressing moderate levels of viral transcripts (below 50% of all detected transcripts) and other cells up to 95%. Additionally, we identified four host genes, *CDH1*, *CORO1C*, *ISCU* and *TMEM119*, whose expression levels strongly correlated with the extent of viral reactivation. Through overexpression and silencing studies, we show that *TMEM119*, a gene with unknown functions, plays an important direct role in promoting KSHV reactivation.

Second, the ability to fixate infected cells or patient-derived materials greatly facilitates the study of virus infected cells outside high-containment BSL-3 facilities that are often not readily available. To demonstrate a potential application of FD-seq in the ongoing COVID-19 pandemic, we applied it to study the infection of OC43, a human betacoronavirus and a close relative of SARS-CoV-2, at the single-cell level. OC43 is a human pathogen that causes the common cold, and we have successfully used to it discover drugs that inhibit SARS-CoV-2 replication *in vitro*^17^. Using FD-seq, we show that instead of the infected cells, bystander cells that express a low level of viral genes are the actual drivers of pro-inflammatory gene expression during coronavirus infection^18^.

Thus, FD-seq is a valuable tool for studying rare cell populations, and provides flexibility in choosing the cell fixation or pathogen inactivation method for researchers.

## RESULTS

### Development of FD-seq for sequencing of PFA fixed single cells

In the standard Drop-seq protocol, single cells are partitioned with uniquely barcoded ceramic beads inside nanoliter droplets in oil, using a microfluidic device^3^. The cells are individually lysed inside the droplets, and their mRNAs are captured by the oligonucleotides on the beads. Next, the droplets are broken to recover the beads, and the beads are extensively washed to remove uncaptured mRNAs. After pooling the beads, the captured mRNAs undergo reverse transcription, exonuclease I digestion of free oligonucleotides on the beads, and whole transcriptome amplification. Finally, the barcoded and amplified cDNAs are tagmented and sequenced.

PFA fixation of cells induces crosslinking between nucleic acids and proteins. To extract RNA from PFA-fixed cells, a cross-link reversal step is therefore required. First, we tested different heating conditions for the cross-link reversal step at bulk level, followed by lysing the uncross-linked cells with Drop-seq lysis buffer, and total RNA extraction using a commercial kit (**Fig. 1a** and **S1**, see Methods). We found that a 1-hour incubation at 56°C in the standard Drop-seq lysis buffer efficiently reversed PFA cross-linking, in agreement with previous literature^5^. The RNA yield was further improved by the addition of proteinase K to the lysis buffer at an optimal concentration of 40 units/mL (**Fig. 1a, Fig. S1a**). Whereas proteinase K treatment did not significantly affect the RNA quality at any of the tested concentrations, consistently resulting in high quality total RNA demonstrated by high (above 8.0) RNA integrity numbers (**Fig. S1b,c**), we observed that increasing the proteinase K concentration above 40 units/mL resulted in a reduction in the overall RNA yield (**Fig. S1a**).

**Figure 1.**
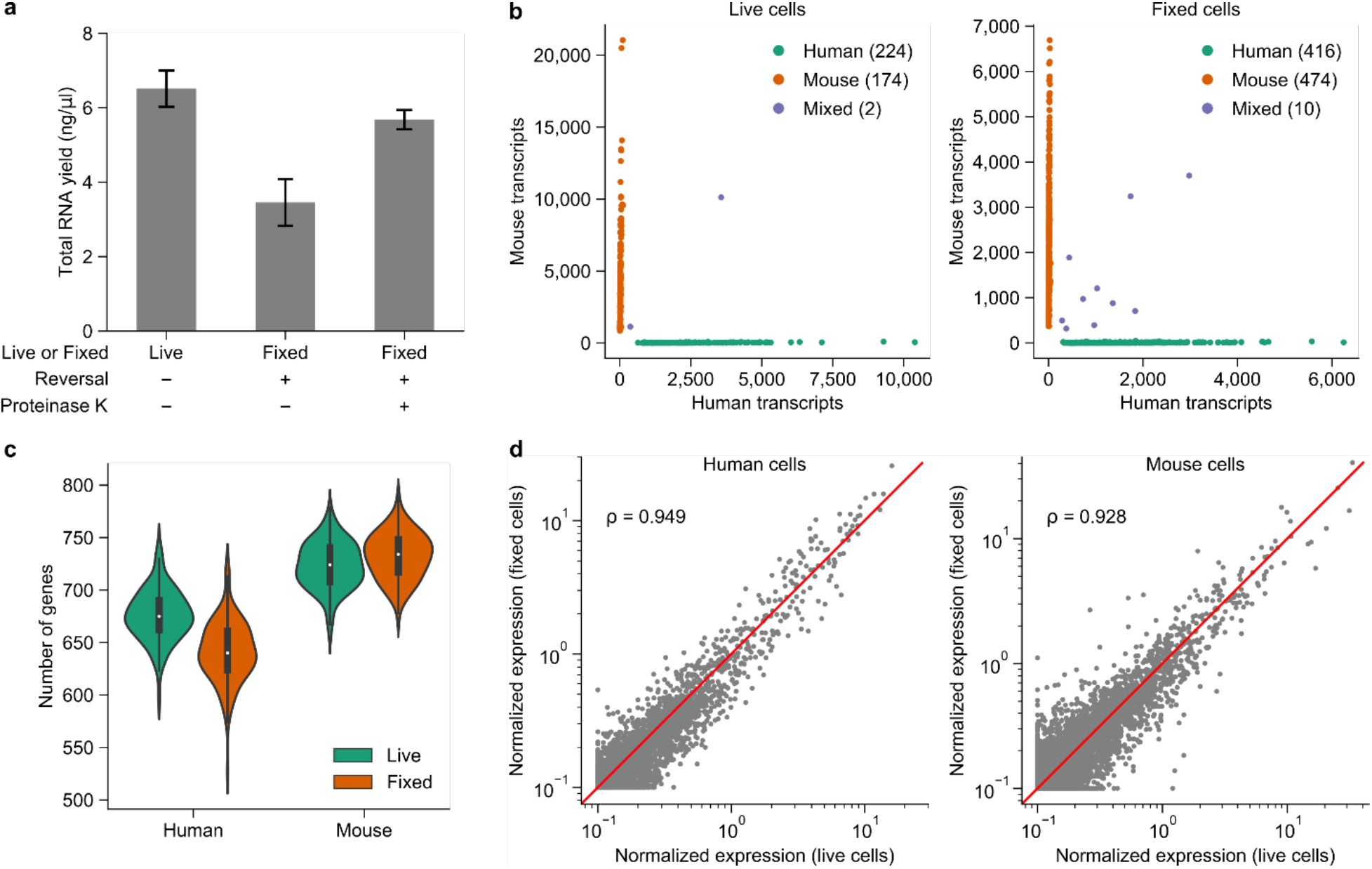
Benchmarking and validation of FD-seq. (**a**) Bar plots showing RNA yield from bulk live cells, and fixed cells that underwent cross-link heat reversal (1 h at 56°C) with or without 40 units/ml of proteinase K. Data is presented as mean ± s.e.m. (**b**) Species-mixing plot of Drop-seq and FD-seq. The multiplet rate for live cells and fixed cells were ~0.5% and ~1%, respectively. For Drop-seq experiment, live human BC3 cells were combined with live mouse 3T3 cells and processed with the standard Drop-seq protocol. For FD-seq experiment, the human and mouse cells were fixed separately, then pooled and processed with the modified FD-seq protocol. (**c**) Violin plots and box plots showing the number of detected genes in live and fixed cells for each species. For this analysis, only cells with at least 1,500 transcripts were considered, and 1,000 transcripts were randomly sampled from each single cell. The white dots inside the violin plots represent the median of the data, the black boxes represent the first and third quartiles, and the black lines represent the values 1.5× the interquartile range beyond the first and third quartiles. (**d**) Comparison of the normalized expression level of each gene between live and fixed cells for each species. Each dot represents a gene, and the red line indicate the line y = x. The plot also shows the Pearson’s correlation ρ of the log-normalized expression level between live and fixed cells.

Next, we compared the extent of inter-droplet RNA contamination between the standard Drop-seq protocol on live cells and the FD-seq method on PFA-fixed cells by performing ‘species-mixing’ experiments. We either used the standard Drop-seq protocol to analyze a 1-to-1 mixture of live BC3 cells, a human PEL cell line, and mouse 3T3 cells, or used FD-seq to analyze BC3 and 3T3 cells that were fixed with 4% PFA and permeabilized with 0.1% Triton-X separately before combining them at a 1-to-1 ratio. The RNA-seq results were aligned to a combined human-mouse reference genome, and the number of human and mouse transcripts per single cell barcode was then calculated. The rate of cell barcodes having both mouse and human transcripts, which indicates cross-droplet contamination and/or multiple cells captured in the same droplet, were very similar and only minimally observed in both live and fixed cell experiments (~0.5% and ~1% for live and fixed cells, respectively, **Fig. 1b**). This indicates that the modifications of FD-seq did not affect the single-cell capture efficiency of Drop-seq.

These experiments also showed that the number of genes detected in fixed cells using was comparable to that of live cells across species (median of 640 genes in fixed cells compared to 675 genes in live cells for human cells, and median of 734 genes compared to 724 genes for mouse cells) (**Fig. 1c**). The relative expression levels of the detected genes were well correlated between live and fixed cells (**Fig. 1d**), even though fewer transcripts were detected in fixed cells than in live cells on average (**Fig. 1b, Fig. S2a**). Finally, live and fixed cells showed a similar percentage of reads mapped to introns and exons (**Fig. S2b**).

Taken together, these results demonstrate that FD-seq is a reliable method for whole transcriptome analysis of PFA-fixed single cells. The performance of single-cell sequencing is maintained with fixed cells when FD-Seq is utilized, and FD-seq shows comparable performance to the standard Drop-seq protocol used for live (unfixed) cells. Reliable sequencing of PFA fixed cells with high transcript recovery can be very useful for single cell studies.

### PFA fixation detects a higher number of genes and transcripts compared to methanol fixation in single cells

We next investigated how the effects of PFA fixation in FD-seq compare to methanol (MeOH) fixation. Methanol fixation has recently been shown to be compatible with Drop-seq^9^ by introducing a suitable rehydration step before droplet generation. Here we fixed A549 cells, a human lung epithelial cell line, with either PFA or methanol, and then analyzed the single-cell transcriptomes by RNA-seq using the FD-seq or Drop-seq protocol, respectively (see Methods).

We found that PFA fixation resulted in a higher number of detected transcripts, or UMIs, (median 8,083 compared to 6,194 transcripts) and genes (median 3,049 compared to 2,856 genes), and a lower percentage of mitochondrial genes (median 6.2% compared to 11.9%) than methanol fixation (**Fig. 2a**). To determine the effect of sequencing depth on the number of UMIs and genes, we randomly sub-sampled the sequencing reads and found that PFA fixation still returned a higher number of UMIs and genes (**Fig. 2b**). The effect of sequencing depth on the number of transcripts is much more pronounced: the difference between the two fixation techniques could reach almost 2,000 UMIs. This could be because a higher proportion of reads in methanol-fixed cells mapped to the intronic and intergenic regions (**Fig. 2c**). Despite this, the average gene expression levels were well correlated between the two fixation methods (Pearson’s correlation coefficient was approximately 0.90, **Fig. 2d**). Lastly, we estimated the technical variance of each method by calculating the gene-level coefficient of variation (CV)^19^. As expected, for both fixation methods, low- and high-abundance genes had higher and lower variance, respectively. In addition, both methods exhibited very similar technical variance. In summary, FD-seq was able to recover more genes and transcripts than methanol fixation, and both PFA- and methanol-fixed cells showed similar average gene expression levels and technical variance.

**Figure 2.**
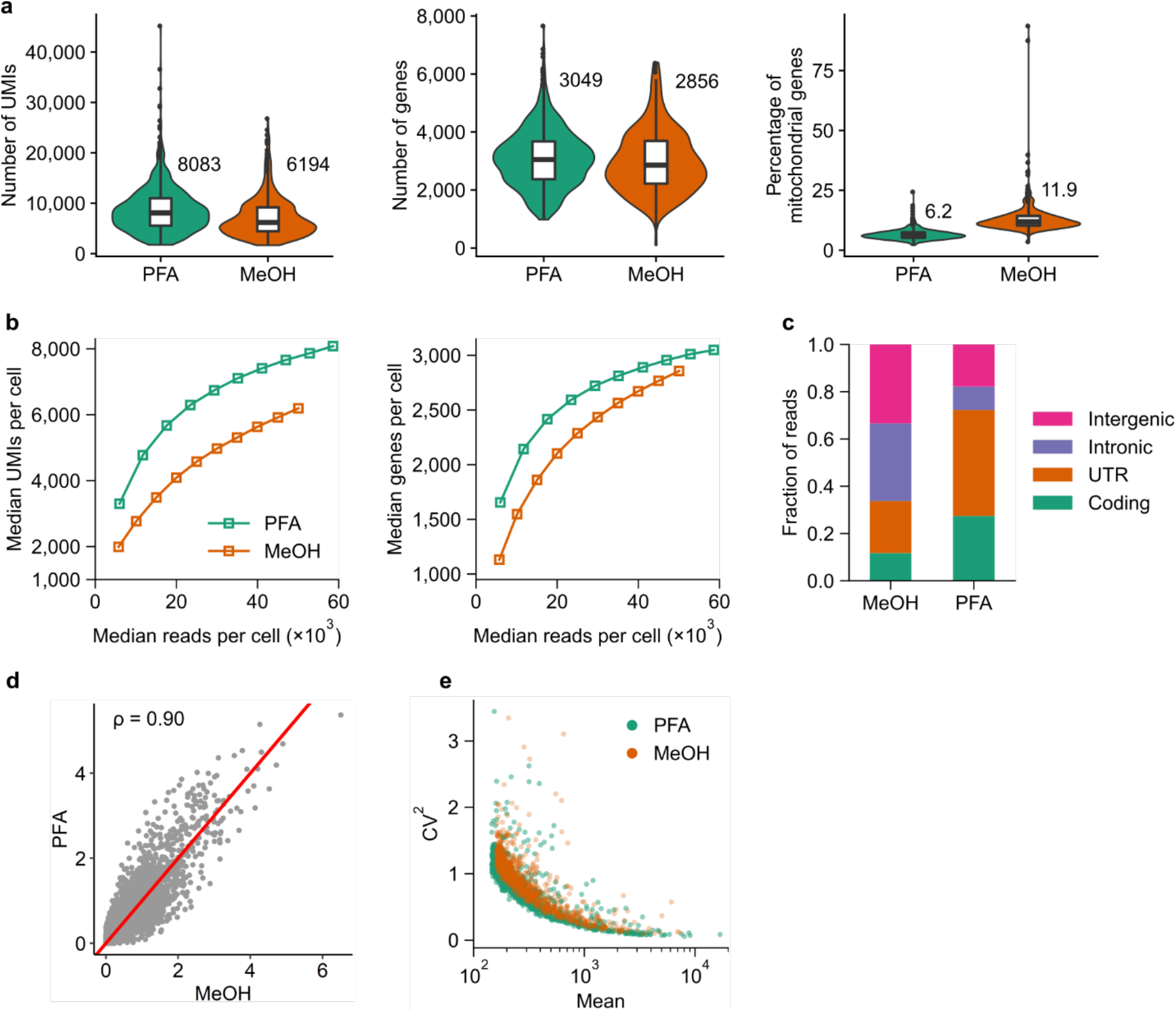
Comparison between PFA and methanol fixation shows higher gene and transcript recovery with FD-seq. (a) Violin and box plots of the number of UMIs, the number of genes and the percentage of mitochondrial genes detected in single A549 cells for each fixation method. The numbers by the violin plot indicate the median values. The middle line inside the box indicates the median, the upper and lower edges of the box indicate the first and third quartiles, and the whiskers extend to 1.5× the interquartile range beyond first and third quartiles. (b) The effects of sequencing depth on the number of detected UMIs and genes per cell. (c) The distribution of mapped reads to different genomic regions. (d) Correlation of the log-normalized average expression level of each gene between the two fixation methods. The plot also shows the Pearson’s correlation coefficient ρ. The red line indicates the y=x line. (e) Technical variance of each gene estimated by the gene’s mean and squared coefficient of variation (CV^2^).

### FD-seq reveals heterogeneity in KSHV reactivated single tumor cells

Having established the validity of FD-seq for the single-cell transcriptome analysis of PFA-fixed cells, we applied FD-seq to test the hypothesis that the heterogeneity in specific host genes contributed to the restricted KSHV reactivation, and to identify those genes.

To identify lytically reactivated cells, we measured expression of the intracellular viral glycoprotein K8.1 that is expressed during lytic reactivation by flow cytometry. As expected, we did not observe any appreciable K8.1 expression in untreated BC3 cells (**Fig. S3a**). However, treatment with NaBut or TPA to induce KSHV reactivation resulted in a small fraction of cells expressing K8.1 (~8% vs ~2.1%, respectively) (**Fig. S3b**). To enrich for reactivated cells, TPA-treated BC3 cells were fixed and sorted by FACS based on the expression level of the K8.1 protein (**Fig. S4**), followed by single-cell transcriptome analysis with FD-seq. We obtained high-quality transcriptome information for 1,035 K8.1+ (reactivated) single cells and 286 K8.1-(latent, non-reactivated) cells. High dimensional clustering and visualization showed clear separation between reactivated and non-reactivated cells (**Fig. 3a**), and analysis of the *K8.1* mRNA level confirmed the enrichment of the K8.1+ cell population of interest (**Fig. 3b**). Moreover, the high proportion of viral transcripts in the K8.1+ population, 69% on average, compared to only 4% viral transcript in the K8.1-population confirmed that the sorted population was indeed composed of reactivated cells (**Fig. 3c,d**).

**Figure 3.**
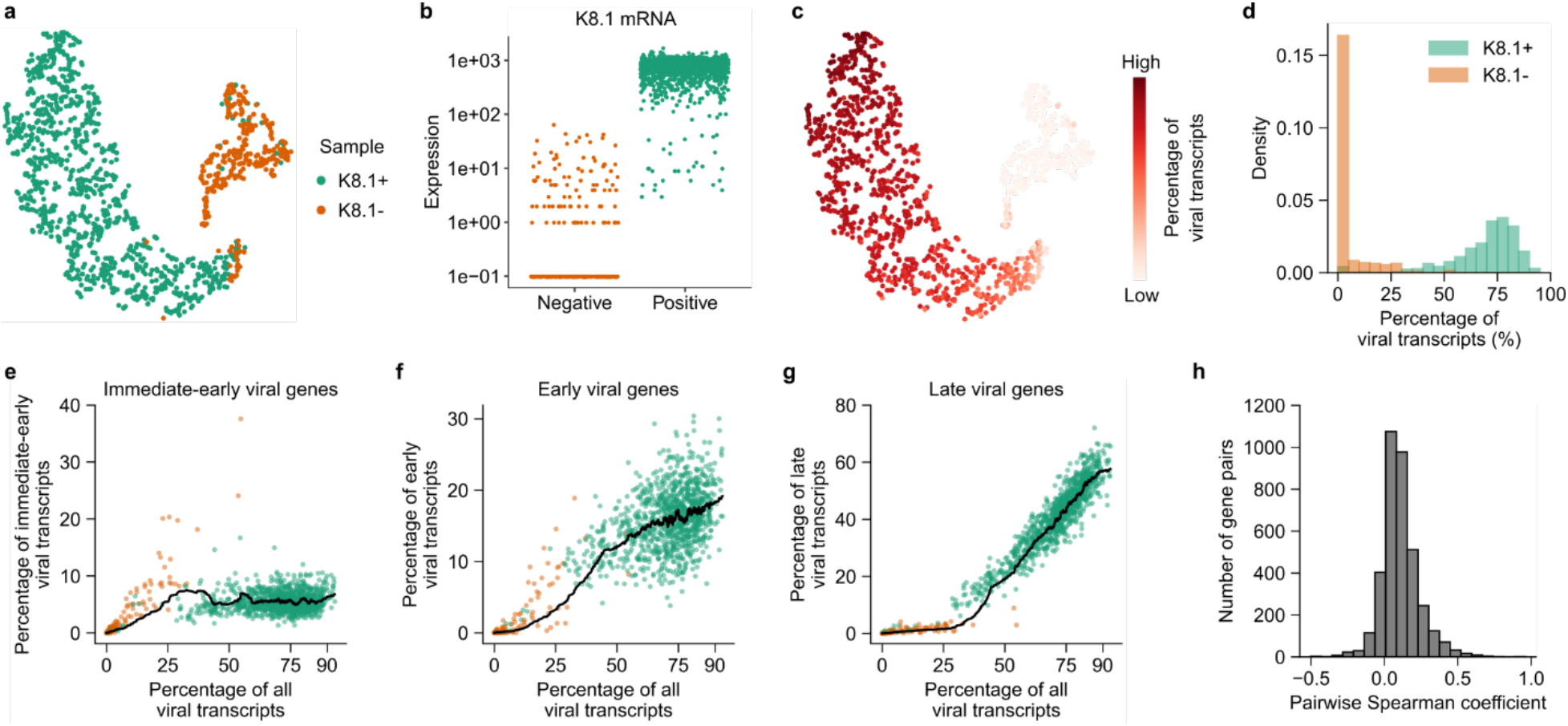
FD-seq reveals heterogeneity in viral reactivation. (**a**) t-SNE plot of K8.1+ (green, lytically reactivated) and K8.1-(red, non-reactivated) BC3 cells as analyzed by FD-seq. (**b**) KSHV K8.1 mRNA levels in the K8.1- and K8.1+ sorted populations. (**c**) t-SNE plot of reactivated and non-reactivated cells as in (a), colored by the percentage of total viral transcripts. (**d**) Histogram showing the percentages of detected transcripts that are from KSHV in K8.1+ and K8.1-BC3 cells. (**e-g**) Change in the percentages of (e) intermediate-early, (f) early and (g) late viral genes as a function of the percentage of total viral transcripts. The black lines indicate the moving averages (50-cell window). (**h**) Histogram of pairwise Spearman correlation coefficients between viral genes.

Herpesvirus reactivation involves the highly regulated, sequential expression of immediate-early, early, and late viral genes^20^ that was recapitulated in our FD-seq results. By ordering the cells by the percentage of viral genes relative to the total transcript content, we found that the relative abundance of immediate-early viral genes increased with total viral gene content, then plateaued after the total viral transcript abundance reached 50% (**Fig. 3e-g**). Early viral transcripts also increased early and monotonically with the abundance of total viral transcripts, without plateauing like the immediate-early genes. On the other hand, late viral transcripts were only detected in cells with more than 25% total viral transcripts, and increased strongly with higher total viral transcript abundance. Thus, FD-seq’s results are in agreement with the expected kinetics of viral gene expression obtained from population-averaged measurements, and suggest that the percentage of viral transcript content is a good indicator of the stage of KSHV reactivation.

Interestingly, we found that the K8.1+ population was highly heterogeneous in viral transcript expression: the relative abundance of viral transcripts varied from below 50% to over 90% (**Fig. 3d**), and the correlation between the expression levels of viral genes was very low, even between viral genes that have the same kinetics (**Fig. 3h** and **Fig. S5a**). This agrees with two recent studies that showed heterogeneity in host cell factor abundance at the single-cell level in herpes simplex virus type 1 (HSV-1) infection^21^, and low correlation in expression of viral genes in a murine gammaherpesvirus infection model^22^. This may partially be caused by the fact that the K8.1+ cells were at different stages of reactivation. Indeed, flow cytometry analysis showed that K8.1 protein abundance within the positive population varied over one order of magnitude (**Fig. S4**). However, this alone could not sufficiently explain the poor correlation between viral genes, because we also observed low correlation coefficients among cells with similar viral transcript abundance that are thought to be in the same stages of reactivation (**Fig. S5b**).

In short, we used FD-seq to successfully characterize KSHV-infected cells undergoing reactivation from latency, revealing the highly heterogeneous nature of this process.

### *TMEM119* facilitates KSHV reactivation

Next, we sought to identify host genes that facilitate KSHV reactivation by looking for differentially expressed host genes that are positively correlated with viral transcript abundance. We only looked for passively correlated host genes in our analysis because the majority of differentially expressed genes were negatively correlated with the relative abundance of viral transcripts (**Fig. S6a**). While this could suggest that their expression inhibits KSHV reactivation^23^ (and therefore cells with lower expression are more prone to reactivate), this type of analysis may be confounded by the high proportion of viral transcripts in the K8.1^+^ population (up to 96%, **Fig. 3d**) that prevents reliable determination of host transcript levels in these cells.

We found four host genes whose expression positively correlated with the abundance of KSHV transcripts: *ISCU*, *CDH1*, *CORO1C* and *TMEM119* (q-value < 10^−100^ for all four genes). We observed that the expression of *ISCU* was relatively low in non-reactivated cells, and increased significantly with the abundance of viral transcripts (**Fig. 4a**). On the other hand, *CDH1*, *CORO1C* and *TMEM119* were mostly undetectable in non-reactivated cells, and increased by 1-2 orders of magnitude in cells expressing high levels of KSHV. We also confirmed the upregulation of *ISCU*, *CORO1C* and *TMEM119* following KSHV reactivation in bulk cell samples by qPCR (**Fig. S6b,c**).

**Figure 4.**
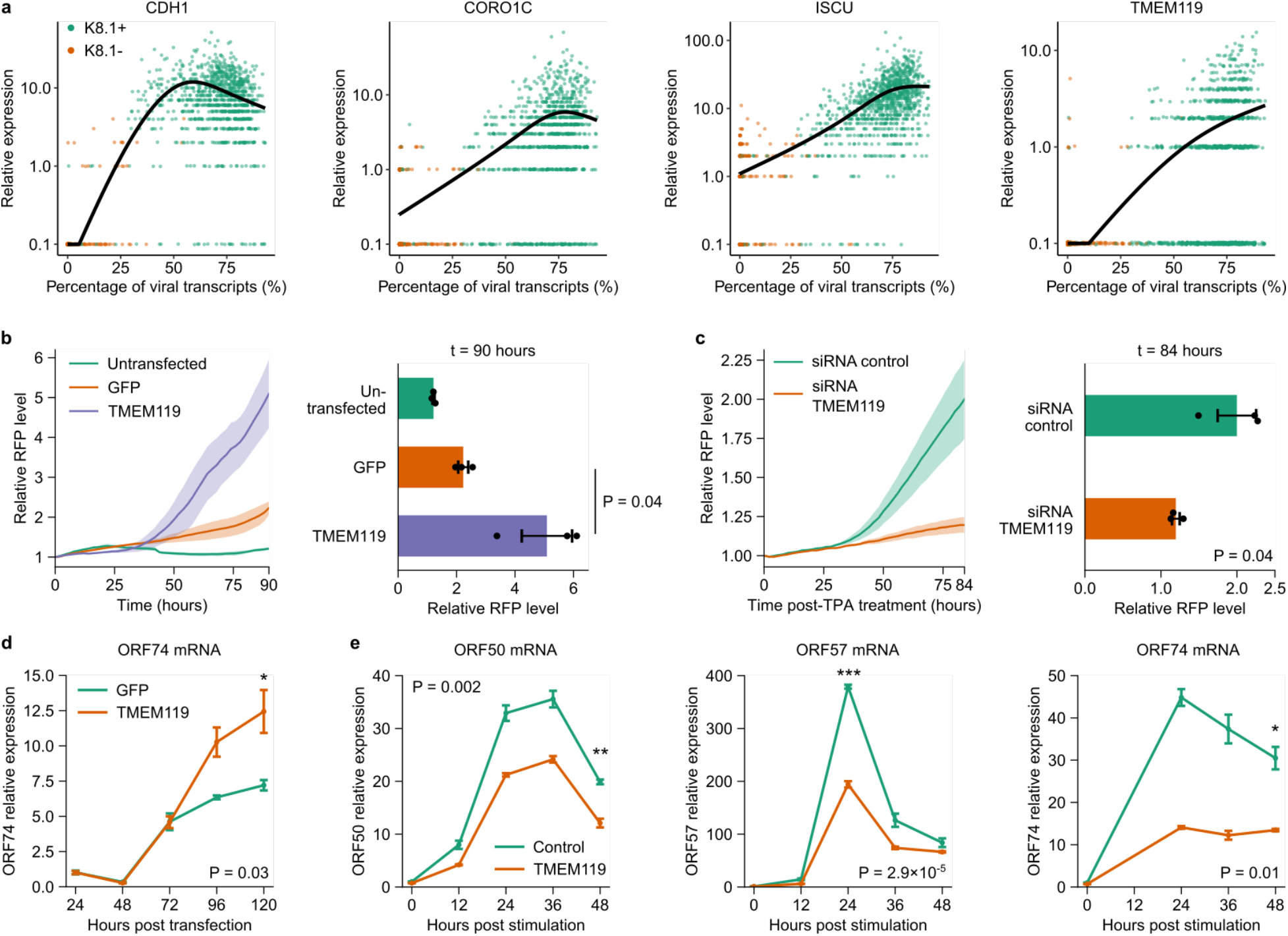
FD-seq identifies *TMEM119* as a host factor that mediates KSHV reactivation. (**a**) Correlation between the percentage of KSHV transcripts and the relative expression of the four host factors *CDH1*, *CORO1C*, *ISCU* and *TMEM119*. Each dots indicate a single cell. (**b**) Live-cell imaging analysis of RFP expression, which is indicative of KSHV reactivation, in untransfected HEK293T.rKSHV219 cells or cells transfected with *TMEM119* or GFP control (left), and end-point quantification of RFP expression at 90 h (right). (**c**) Live-cell imaging analysis of RFP expression in HEK293T.rKSHV219 cells transfected with control or TMEM119-targeting siRNAs for 48 h, followed by treatment with 2 ng/mL TPA (left), and end-point quantification of RFP expression at 84 h (right). The ribbons in (b,c) indicate the s.e.m. of the data. P-values: one-sided Welch’s t-test. (**d**) Time-course qRT-PCR analysis of KSHV *ORF74* abundance in HEK293T.rKSHV219 cells transfected with *TMEM119* or GFP control. (**e**) Time-course qRT-PCR analysis of KSHV *ORF50*, *ORF57*, and *ORF74* levels in HEK293T.rKSHV219 cells transfected with control or *TMEM119*-targeting siRNAs. (b-e) Data are represented as mean ± s.e.m. (n=3 biological replicates). (d,e) *P < 0.05, **P < 0.01, ***P < 0.001 (one-sided Welch’s t-test).

To determine whether the strong correlation between these host transcripts and KSHV gene expression means that these genes modulate KSHV reactivation induction and/or efficiency, we next tested the effect of their overexpression on KSHV reactivation. To this end, we used live-cell imaging to monitor KSHV reactivation in HEK293T.rKSHV219 cells (which are HEK293T cells latently infected with the recombinant KSHV.219 virus strain that constitutively expresses GFP and further encodes RFP under control of the viral lytic PAN promoter^24^) following exogenous expression of *CDH1*, *CORO1C*, *ISCU*, or *TMEM119*. Whereas overexpression of *CDH1*, *CORO1C*, or *ISCU* had no significant effect on KSHV reactivation efficiency, we observed a clear increase in RFP expression, which is indicative of viral reactivation, upon ectopic expression of *TMEM119* relative to a GFP-encoding control vector (**Fig. 4b, Fig. S7a-c**). Conversely, silencing of endogenous *TMEM119* significantly reduced the level of TPA-induced KSHV reactivation (**Fig. 4c, Fig S7b,c**). These results were corroborated by qRT-qPCR analysis of KSHV lytic gene expression, which showed that *TMEM119* overexpression enhanced KSHV *Orf74* expression compared to the *GFP* control (**Fig. 4d**), while *TMEM119* silencing led to a reduction in KSHV *Orf50, Orf57, and Orf74* expression (**Fig. 4e**). Together, these results show that *TMEM119* positively modulates KSHV reactivation efficiency.

Not much is known about the function of *TMEM119*, which has been previously characterized as a marker for microglia in the human brain^25^. To determine the potential mechanism by which *TMEM119* enhances KSHV reactivation, we performed bulk transcriptome analysis of HEK293T cells with and without ectopic expression of *TMEM119*. We found that in HEK293T cells, 202 genes were significantly upregulated upon *TMEM119* overexpression (**Fig. S8a** and **Supplementary File 1**). KEGG pathway analysis^26^ showed that these genes were highly enriched for the MAPK signaling pathway (**Fig. S8b**). This finding is interesting, as previous works have implicated the MAPK pathway in KSHV reactivation^27,28^. Together with this, our data suggests that *TMEM119* promotes viral reactivation via activation of the MAPK signaling pathway.

### FD-seq reveals a pro-inflammatory subpopulation in OC43 infected cells

We next applied FD-seq to study the infection of the betacoronavirus OC43 in single human lung cells. OC43 is a human pathogen that causes the common cold, and is a close relative of SARS-CoV-2. We have recently shown that many drugs that inhibit the replication of OC43 also inhibit SARS-CoV-2 replication *in vitro*^17^, suggesting that OC43 is a good model system for SARS-CoV-2 infection.

First, we infected A549 cells with OC43 at a multiplicity of infection (MOI) of 1, fixed the cells with PFA, and performed FD-seq on mock- and OC43-infected cells. We obtained 1,167 and 1,924 high-quality single cells from mock-infected and OC43-infected cells, respectively. After regressing out the effects of cell cycle variations (**Fig. S9a, b**), high dimensional clustering yielded three main clusters of cells (**Fig. 5a**). Cluster 0 mainly contained the mock-infected cell, while clusters 1 and 2 mainly contained OC43-infected cells (**Fig. 5b**). At an MOI of 1, more than 70% of the cells showed some level of viral gene expression (**Fig. 5c**). Interestingly, this infected population consisted of two subpopulations based, clearly separated by the number of viral transcripts detected: a larger subpopulation containing a low number of viral transcripts (below 100 UMIs, median 2 UMIs) and a smaller subpopulation containing a much higher number of viral transcripts (above 100 UMIs, median 428 UMIs) (**Fig. 5d**). This small subpopulation of highly infected cells corresponded to cluster 2 (**Fig. 5e** and **Fig. S9c, d**).

**Figure 5.**
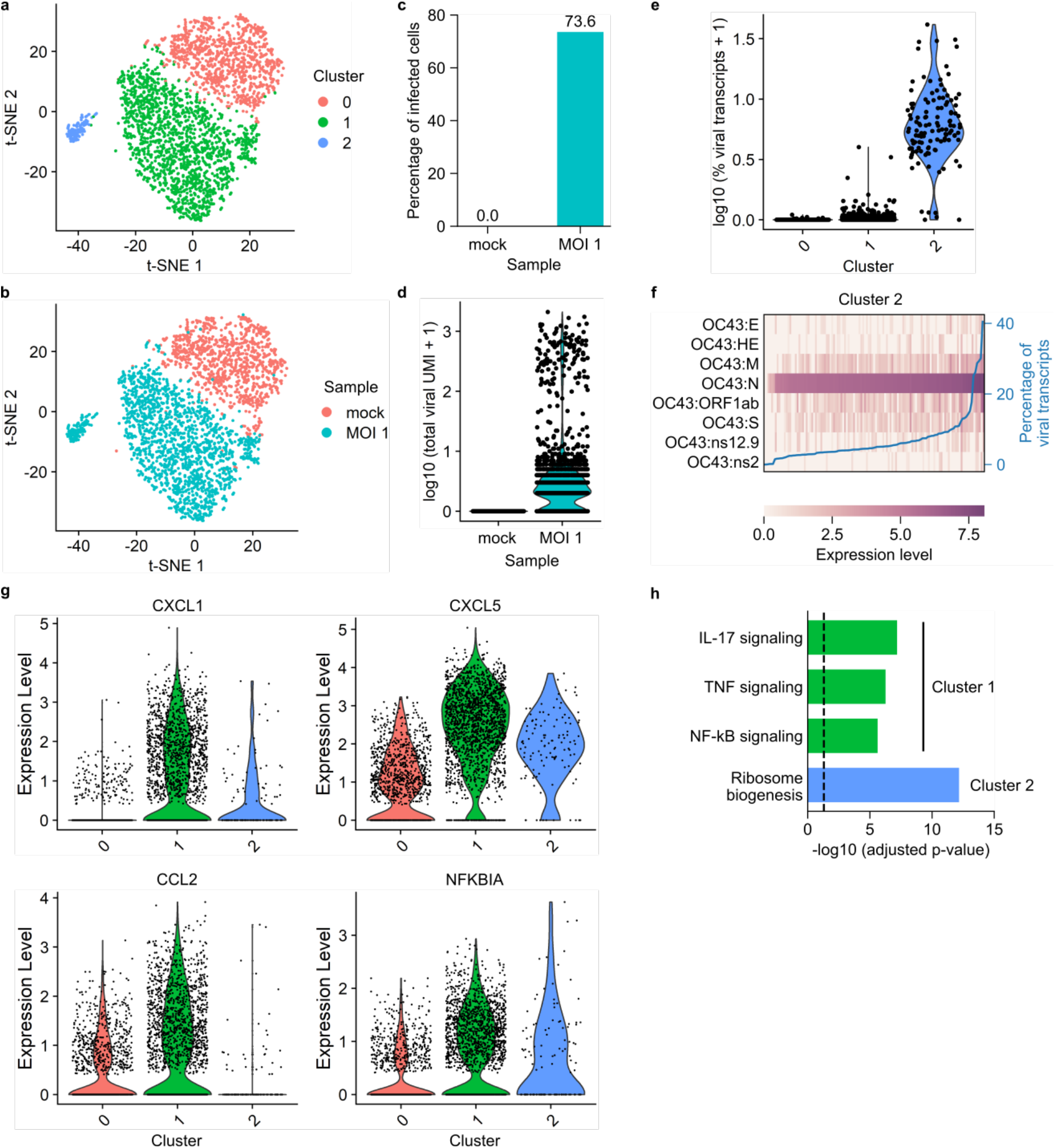
FD-seq reveals pro-inflammatory bystander cells after coronavirus OC43 infection. (**a-b)** t-SNE plots with cells colored by (a) cluster identity or (b) sample type. (**c)** Bar plot showing the percentage of infected cells (defined as cells that expressed at least 1 detected viral transcript) of mock infected and MOI 1 sample. (**d)** Violin plot showing distribution of total viral transcript counts. (**e)** Violin plot showing the distribution of the percentage of total viral transcript by cluster identity. (**f)** Heatmap showing the relative expression level of each viral gene of each single cell in cluster 2. The blue line shows each cell’s percentage of total viral transcripts. (**g)** Violin plots showing the expression of four representative immune-related genes that are upregulated in cluster 0. (**h)** Bar plot showing the adjusted P-values of upregulated KEGG pathways in clusters 1 and 2. The vertical dashed line indicates P-value = 0.05.

Next, we sought to investigate the expression profile of OC43 viral genes as a function of total viral gene level within cluster 2’s cells. Gene *N* (which encodes the viral nucleocapsid protein^29^) was uniformly highly expressed (**Fig. 5f**). The expression of the remaining viral genes was more heterogeneous, with *ORF1ab*, *M* (membrane gene) and *S* (spike gene) being more highly expressed overall. These patterns of viral gene expression level were also true when looking at all infected cells (**Fig. S10**).

Cluster 1 consisted of cells that were exposed to the virus but failed to express high levels of viral genes (**Fig. 5d** and **Fig. S8c, d**), and were thus either abortively-infected or uninfected bystander cells. We found that these bystander cells were enriched in pro-inflammatory genes, such as *CXCL1*, *CXCL5*, *CCL2* and *NFKBIA* (**Fig. 5g, Fig. S11**, and **Supplementary File 2**). *CXCL1* and *CXCL5* encode the protein ligands of the CXC chemokine receptor 2 (CXCR2), and are crucial to neutrophil recruitment^30^. *CCL2* encodes a different chemokine that is responsible for monocyte recruitment from the bone marrow^31^. *NFKBIA* encodes the inhibitor IκBα to the transcription factor NF-κB, which is an important regulator of the immune response^32^. KEGG pathway analysis showed that cluster 1 was enriched for three main pro-inflammatory pathways: TNF, IL-17 and NF-κB signaling pathways (**Fig. 5h**). On the other hand, the ribosome biogenesis pathway is enriched in the upregulated genes of cluster 2, suggesting an increased need for protein biogenesis during OC43 replication.

In short, FD-seq revealed that the majority of cells did not express a high level of viral genes after exposure to OC43, and that these cells upregulated pro-inflammatory genes. These findings are in agreement with our previous characterization of HSV-1 infection at the single-cell level^21^, which showed that most HSV-1 infected cells did not express high levels of viral genes, and that these abortively-infected cells were the main producers of antiviral gene transcripts and the initiators of the type I interferon signaling.

## DISCUSSION

Here we present FD-seq, a novel method for high-throughput droplet-based RNA sequencing of PFA-fixed single cells. FD-seq is particularly useful for sequencing rare subpopulations of cells that require intracellular staining and FACS-enrichment, and for rendering infectious samples safe for handling. Drop-seq has only been shown to work with methanol fixation^9^, and we demonstrated that FD-seq performed better than methanol fixation, with the former detecting more genes and transcripts. FD-seq will increase the flexibility for researchers in using high throughput scRNA-seq, because PFA fixation has also shown to provide a better signal-to-background ratio for certain intracellular staining targets^11,12^, and because PFA is a very commonly used alternative to methanol fixation.

We applied FD-seq to study the process of KSHV lytic reactivation at the single-cell level, and found that reactivation is very heterogeneous, and that the expression levels of viral genes correlated poorly with one another. We found four host factors, namely *CDH1*, *CORO1C*, *ISCU* and *TMEM119,* to be significantly positively correlated with viral reactivation. Using live-cell imaging and time-course study, we found *TMEM119* to have the most pronounced effect among the four genes in enhancing the degree of viral reactivation. Bulk RNA-seq suggested that this effect of *TMEM119* could be mediated through the MAPK signaling pathway, in agreement with previous studies ^27,28^. More studies are required to investigate in details the mechanisms by which *TMEM119* affects the MAPK signaling pathway and modulates the increase in KSHV reactivation.

Furthermore, we demonstrated a potential application of FD-seq in the COVID-19 pandemic, by using FD-seq to study the host response of human lung cells to OC43, a betacoronavirus, infection, and the subsequent expression of viral genes at the single-cell level. First, we infected the cells with the virus, then fixed with PFA, thereby inactivating the virus for safer sample handling, before analyzing the cells with FD-seq. We showed that most OC43-infected cells were unable to support high level of viral gene expression, and instead upregulated pro-inflammatory signaling pathways. Therefore, these lowly-infected cells are the main producers of pro-inflammatory gene products, such as cytokines, during OC43 infection *in vitro*. It would be very interesting to examine whether a similar phenomenon is observed for SARS-CoV-2 infected cells, both *in vitro* and in patient samples. FD-seq would allow researchers to inactivate this infectious virus with PFA so that they can safely study it in BSL-2 facilities.

Taken together, these biological application of FD-seq show that this method is a valuable tool for integrating protein activity with transcriptome information. Transcription factor expression levels and phosphorylation can be integrated with whole transcriptome analysis by combining intracellular protein staining with FD-seq. Furthermore, FD-seq is compatible with RNA velocity analysis, which relies on detection of unspliced mRNA molecules^33^ (**Fig. S12**). In future studies, FD-seq could serve as a basis for developing a method for sequencing formalin-fixed paraffin-embedded (FFPE) tissues, due to the similarity between PFA fixation and FFPE. Such an achievement of which would enable high-throughput single-cell sequencing of readily available samples in tissue banks.

## ACKNOWLEDGEMENTS

We thank Heather Eckart for providing advice on Drop-seq. This study was supported in part by U.S. National Institutes of Health (NIH) grants (R21 AI118509 and R01 AI087846 to M.U.G.) and P. G. Allen Distinguished Investigator Award, NIH grants GM127527 and GM128042 to S.T.

## MATERIALS AND METHODS

### Microfluidic device fabrication

The microfluidic device was fabricated based on the AutoCAD file provided in Macosko et al.^3^ The mold was made from SU8-3050 photoresist of 100 μm height. Then, polydimethylsiloxane (PDMS, Momentive RTV615) was cast on the mold to form the microfluidic devices. Plasma bonding was used to bond the PDMS device to a glass slide, and the microfluidic channels were coated with Aquapel.

### Cell culture

HEK293T (ATCC) cells were maintained in Dulbecco’s Modified Eagle’s Medium (DMEM) supplemented with 10% (v/v) fetal bovine serum (FBS), 2 mM GlutaMAX (Gibco), and 1% (v/v) penicillin-streptomycin (P/S, Gibco). HEK293T.rKSHV.219 cells were generated by infecting HEK293T cells with rKSHV.219^24^, and selecting with 1 μg/mL puromycin. The KSHV-positive human PEL cell line BC3 was cultured in Roswell Park Memorial Institute (RPMI) supplemented with 20% (v/v) FBS, 2 mM GlutaMAX, and 1% (v/v) P/S. A549 cells (ATCC) or A549 cells with H2B-Ruby fusion were maintained in DMEM supplemented with 10% FBS, without antibiotics. The 3T3 cells (p65^−/−^ 3T3 mouse embryonic fibroblast cells expressing p65-DsRed and H2B-GFP nucleus marker^32^) were cultured in DMEM supplemented with 10% (v/v) fetal bovine calf serum (HyClone), 1% (v/v) GlutaMAX and 1x P/S.

### Optimization of RNA extraction condition for fixed and permeabilized cells

BC3 cells were harvested, centrifuged at 300g for 3 min to remove the cell media, and washed once with PBS and 1% BSA (molecular biology grade, Gemini Bio-Products). To fix the cells, 4% PFA in PBS (Santa Cruz Biotechnology) was added to the cells and incubated at room temperature for 15 minutes. Next, the paraformaldehyde was discarded, and the cells was washed once with wash buffer (PBS with 1% BSA and 40 units/ml of RNase inhibitor, murine (NEB)). To permeabilize the cells, 0.1% Triton X-100 (molecular biology grade, Acros Organics) diluted in the wash buffer was added to the cells, and incubated at room temperature for 15 minutes. Then, after adding the wash buffer, cells were pelleted, the supernatant was discarded, and the cells were wash once more with the wash buffer. To mimic the condition of antibody staining, cells were resuspended in wash buffer and incubated on ice for 1 hour, washed twice with the wash buffer and finally kept on ice in the wash buffer before RNA extraction.

To serve as a positive control, RNA from bulk live cells was extracted using the RNeasy Plus Mini Kit (Qiagen). Cells were first suspended in a combination of the standard Drop-seq lysis buffer with PBS at equal volume ratio (to make the lysis buffer’s final concentration the same as that in nanoliter droplets), incubated at room temperature for 10 minutes, and placed on ice for at least 5 minutes. Next, the cell lysate was combined with 350 μl of RLT plus buffer from the RNeasy kit, and processed according to the manufacturer’s protocol.

To extract RNA from bulk fixed cells, fixed and permeabilized cells (as described above) were suspended in a combination of Drop-seq lysis buffer spiked in with various concentrations of proteinase K (NEB), and PBS at equal volume ratio. Next, the sample was at 56°C for 1 hour, left at room temperature for 10 minutes, and placed on ice for at least 5 minutes. Finally, the cell lysate was combined with 350 μl of RLT plus buffer, and processed according to the RNeasy Plus Mini Kit’s protocol.

### Treatment of BC3 cells, flow cytometry and FACS-sorting of reactivated cells

BC3 cells were mock-treated or treated with a final concentration of 20 ng/mL tetradecanoyl phorbol acetate (TPA, Millipore Sigma) for 36 hours to induce KSHV reactivation. Cells were pelleted by centrifugation at 300g for 3 min and fixed with 4% PFA in PBS for 15 min at room temperature followed by permeabilization with 0.1% (v/v) Triton X-100 in PBS with 1% (w/v) BSA for 15 min at room temperature. After washing and blocking the cells in PBS supplemented with 4% BSA, staining was performed with an anti-KSHV K8.1 antibody (sc-65446, Santa Cruz, 1:50 dilution) and an Alexa Fluor 488-conjugated goat-anti-mouse secondary antibody (A10667, Life Technologies, 1:4000 dilution) in PBS/1% BSA supplemented with RNase inhibitor, murine (NEB). K8.1-positive and negative cell populations were analyzed on an LSRFortessa flow cytometer (BD Biosciences) and/or sorted on a FACSAriaIIIu cell sorter (BD Biosciences).

### Infection of A549 cells

OC43 (ATCC) were grown and titrated on A549 cells. For FD-seq experiments, A549 cells were seeded in 6-well plates and infected with OC43 at an MOI of 1 the following day. Cells were incubated at 33C for 4 days and were subsequently harvested for FD-seq.

### FD-seq protocol

The protocol for FD-seq is based on the McCaroll lab’s online Drop-seq protocol^3^. In summary, there are only two significant differences: 40 units/ml proteinase K was added to the Drop-seq lysis, and a heating step was added between the droplet generation and droplet breakage steps. First, the barcoded beads were suspended at 120,000 beads/ml in a modified Drop-seq lysis buffer: 200 mM Tris pH 7.5, 6% Ficoll type 400, 0.2% Sarkosyl, 20 mM EDTA, 50 mM DTT and 40 units/ml proteinase K (NEB). Cells were suspended in PBS with 0.01% BSA at 100,000 cells/ml.

For droplet generation, the cells, beads and oil were injected at flow rates of 3, 3 and 12 ml/h, respectively, using syringe pumps. After droplet generation, the droplets were heated on a heat block at 56°C for 1 hour to reverse the PFA cross-links, then incubated at room temperature for 10 minutes and kept on ice for at least 5 minutes. After this, the droplets were broken, the beads were collected, washed, and subjected to reverse transcription and exonuclease I digestion as per standard Drop-seq protocol.

For whole transcriptome amplification (WTA), the beads were distributed into a 96-well plate such that each well contained 5,000 beads. A 50 ul PCR reaction was set up in each well (1X KAPA HiFi Hotstart Readymix, 0.8 uM TSO_PCR primer) using the following thermal cycle program: 95C 3 min; 4 cycles of 98C 20s, 65C 45s, 72C 3 min; 10 cycles of 98C 20s, 67C 20s, 72C 3 min; 72C 5 min; 4C hold. After PCR finishes, the WTA products were cleaned up with 0.6X Ampure XP beads (Beckman Coulter), pooled and quantified with TapeStation (Agilent).

To prepare the sequencing library, 450pg of WTA products were used for each tagmentation reaction (Nextera XT DNA Library Prep Kit, Illumina). First, the WTA products were incubated with the tagment enzyme at 55C for 5 min. Next, the neutralization buffer was added, and the sample was incubated at room temperature for 5 min, in order to neutralize the enzyme. After this, the Nextera PCR Master Mix and custom primers P5-TSO_Hybrid and Nextera_N70X were added, and the sample was thermocycled as following: 95C 30s; 12 cycles of 95C 10s, 55C 30s, 72C 30s; 72C 5 min; 4C hold. Finally, the tagmentation products were cleaned up with 0.6X Ampure XP beads and quantified with TapeStation. The sequences of the primers are given in Table S1.

### Methanol fixation experiment

A549 cells were harvested and split into 2 samples of approximately 1 million cells each, one for PFA fixation and FD-seq, and one for methanol fixation and Drop-seq. PFA fixation was performed as above. For methanol fixation^9^, cells were washed once with 1 ml of 1% BSA, then resuspended in 200 ul of ice-cold PBS. Next, ice-cold 100% methanol was added dropwise into the cells while gently vortexing the tube, and the cells were incubated on ice for 15 minutes. The cells were then rehydrated by centrifuging the cells at 1000g for 5 minutes, then discarding the methanol and washing twice with 1 ml of 0.01% BSA. The rehydrated cells were then counted and processed according to the standard Drop-seq protocol.

### Plasmids and transfections

Expression plasmids encoding human *CDH1*, *TMEM119*, *ISCU* and *CORO1C*, as well as KSHV *ORF50* and nls-eGFP under control of the human EF1α promoter in the pLV backbone were ordered from VectorBuilder. All sequences were verified by Sanger sequencing. HEK293T.rKSHV219 cells were reverse transfected with 0.8 μg plasmid DNA using Lipofectamine2000 (Life Technologies) following the manufacturer’s instructions and seeded in 12 or 24-well plates for analysis by live-cell imaging or qRT-PCR as indicated. For silencing experiments, HEK293T.rKSHV219 cells were reverse transfected with 40 nM siRNA targeting *TMEM119* (Dharmacon siGenome SMARTpool M-018636-01-0005), *CDH1* (M-003877-02-0005), *CORO1C* (M-017331-00-0005), ISCU (M-012837-03-0005), or non-targeting control (D-001206-14-05) in 12 or 24-well plates using Lipofectamine RNAiMAX reagent (Life Technologies) following the manufacturer’s protocol. After 48 hours, the cells were treated with 2 ng/mL TPA as indicated and KSHV reactivation efficiency was assessed by live cell imaging or qRT-PCR.

### Reverse transcription and real-time PCR (RT-qPCR)

Total RNA was extracted by using the HP Total RNA Kit (OMEGA Bio-Tek) using manufacturer’s instructions. RT-qPCR was performed using either a one-step or a two-step protocol. For the one-step protocol, RT-qPCR were performed with equal amounts (25-500 ng) of RNA using the SuperScript III Platinum One-Step qRT-PCR kit with ROX (Thermo Fisher Scientific) on a 7500 Fast Real-Time PCR Machine (Applied Biosystems). Premixed master mixes containing TaqMan primers and probes for each individual gene were purchased from IDT (*GAPDH*, *TMEM119*, *CORO1C*, *ISCU*, and *CDH1*) or Applied Biosystems (18S). For the two-step protocol, reverse transcription was performed using the ProtoScript II First Strand cDNA Synthesis Kit (NEB) according to the manufacturer’s protocol. The kit’s oligo-dT primer was used as the primer in this step. Next, the RT products were diluted by 20-fold in water, and used as the input for qPCR (Luna Universal qPCR Master Mix, NEB). The primers’ sequences are given in Table S2. Relative expression level of each target gene was calculated by normalizing for GAPDH or 18S levels using the Comparative Ct Method (∆∆Ct Method) and presented relative to the control sample.

### Live-cell imaging experiments

HEK293T.rKSHV were seeded in 24-well plates and transfected with plasmids for over-expression or siRNA for knock-down experiments. Cells were then imaged on a Nikon Ti-Eclipse containing an environmental chamber (37C, 100% humidity, 5% CO2). Images were acquired every 4 hours for 4 days.

### Bulk RNA-seq

Total RNA was extracted from cells with the HP Total RNA Kit as above and quantified using NanoDrop. mRNA was isolated from 100 ng of total RNA per sample using oligo dT magnetic beads (NEB) according to the manufacturer’s protocol. The isolated mRNA was then used to prepare sequencing libraries using NEBNext Ultra II Non-directional RNA Library Prep Kit for Illumina and NEBNext Multiplex Oligos for Illumina (Index sets 1 and 2) (NEB). Next, the sequencing libraries were quantified using High Sensitivity Fragment Analyzer (Agilent), normalized to 30 nM, pooled at equimolar, and sequenced with NextSeq High-output kit (single-read 75 bp).

### Sequencing and alignment

Drop-seq and FD-seq libraries were sequenced on a NextSeq 550 machine with the following read distribution: 20 bp for read 1, 60 bp for read 2, and 8 bp for index 1 read. The custom read 1 sequencing primer Read1CustomSeqB was used in place of Illumina’s read 1 primer (Table S1). Alignment was performed using Picard version 2.21.8 (https://github.com/broadinstitute/picard) and Drop-seq tools version 1.13 or version 2.3. The valid cell barcodes were chosen using the “knee plot” method. Bulk RNA-seq libraries were sequenced on a NextSeq 550 machine with the following read distribution: 75 bp for read 1, and 6 bp for index 1 read. Alignment was performed using STAR aligner (version 2.7.3a) and counted using featureCounts. Alignment was performed to a concatenated version of the human genome GRCh38 and OC43 genome NC_006213.1^29^.

### Read subsampling and read mapping

In the FD-seq and methanol fixation comparison experiment, we used samtools view command^34^ to subsample the output BAM file from Drop-seq tools version 2.3 DetectBeadSynthesisErrors command. To calculate the proportion of reads mapped to different genomic regions, we used Picard CollectRnaSeqMetrics command on the output BAM file directly from the STAR aligner. The refFlat file for the GRCh38 reference genome was downloaded from http://hgdownload.cse.ucsc.edu/goldenPath/hg38/database/.

### scRNA-seq data analysis

Data analysis was done using R packages Monocle 2^35–37^q and Seurat 3^38,39^. Data visualization was done using R and Python. For species mixing data, a cell is classified as human or mouse if more than 90% of the detected transcripts are mapped to human or mouse, respectively. dropEst pipeline^40^ was used with argument –L eiEIBA to annotate reads mapping to exon, intron or exon/intron spanning regions.

For the FD-seq and methanol fixation comparison experiment, we first discarded cells with lower than 1,000 UMIs and genes detected in fewer than 5 cells. We then used Seurat 3 to log-normalize and scale the data. To calculate the technical variance, we normalized the UMI count by converting it to UMIs per million, then randomly chose 400 cells from each fixation method and used the 1,000 most highly expressed non-mitochondrial genes to calculate the mean and squared coefficient of variation.

For the KSHV reactivation experiment, we first discarded cells with more than 3,000 genes detected, and cells with fewer than 1,000 UMIs or more than 10,000 UMIs. Then we discarded genes detected in fewer than 5 cells. To find the relative expression of each gene, we first normalized the cells using Monocle 2. To cluster the cells, we first performed dimension reduction with t-SNE using the first 4 principal components, then clustered the cells by setting rho_threshold = 17 and delta_threshold = 11. We then set the percentage of viral transcripts as the pseudotime parameter, and performed a “pseudotime analysis” in order to find genes correlated with the abundance of viral transcripts.

For the OC43 infection experiment, we discarded cells with fewer than 1,000 UMIs or more than 20,000 UMIs. We then discarded genes detected in fewer than 5 cells. We then used Seurat 3 to remove the cell cycle effects, normalize and scale the data. To cluster the cells, we used the first 10 principal components for the FindNeighbors and RunTSNE functions, and resolution=0.2 for the FindClusters function. To find the cluster markers, we used min.pct=0.25 and logfc.threshold=0.25 for the FindAllMarkers function.

### Bulk RNA-seq data analysis

Bulk RNA-seq data was analyzed using the R package DESeq2^41^. To find the enriched KEGG pathways, we uploaded the list of genes that were upregulated in TMEM119 overexpression versus GFP overexpression (log2 fold change > 1, FDR < 0.05) to g:Profiler^26^.

### RNA velocity analysis

The output files of Drop-seq tools DetectBeadSynthesisErrors function were processed with the dropEst pipeline^40^ to tag spliced and unspliced transcripts, and the results were analyzed with Python velocyto package^33^.

### Data availability

The raw data, metadata and count data are deposited in NBCI’s Gene Expression Omnibus (accession number GSE156988).

