## Supplementary Figures for "Fixed single-cell RNA sequencing for understanding virus infection and host response"

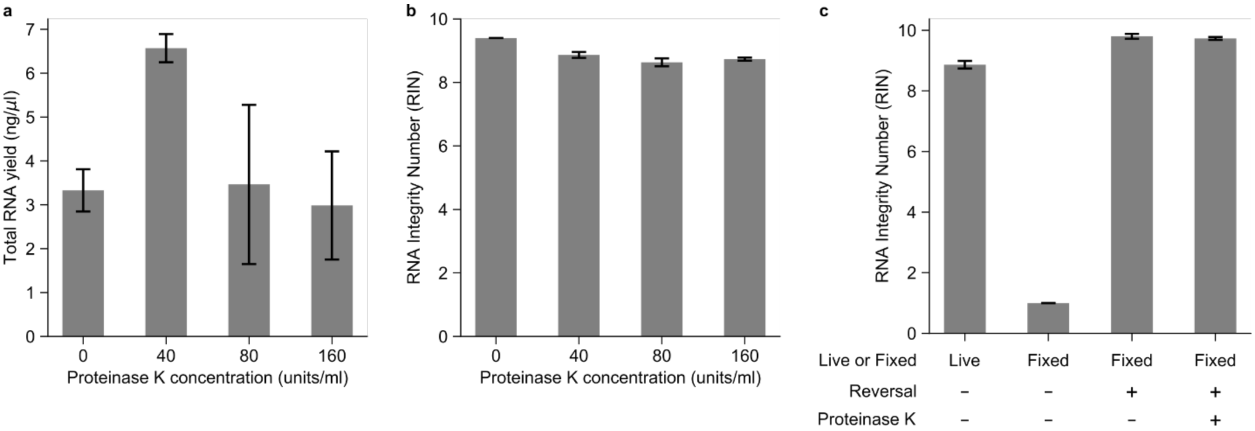


**Figure S1. Optimization of total RNA extraction from bulk fixed cells. (a, b)** Total RNA yield (a) and RNA integrity number (RIN) (b) at different proteinase K concentrations. **(c)** RINs of RNA extracted from fresh live cells, of RNA extracted from fixed cells without heat reversal and without using proteinase K, and of RNA extracted from fixed cells after heat reversal, and with and without using proteinase K. Data is presented as mean ± standard deviation.

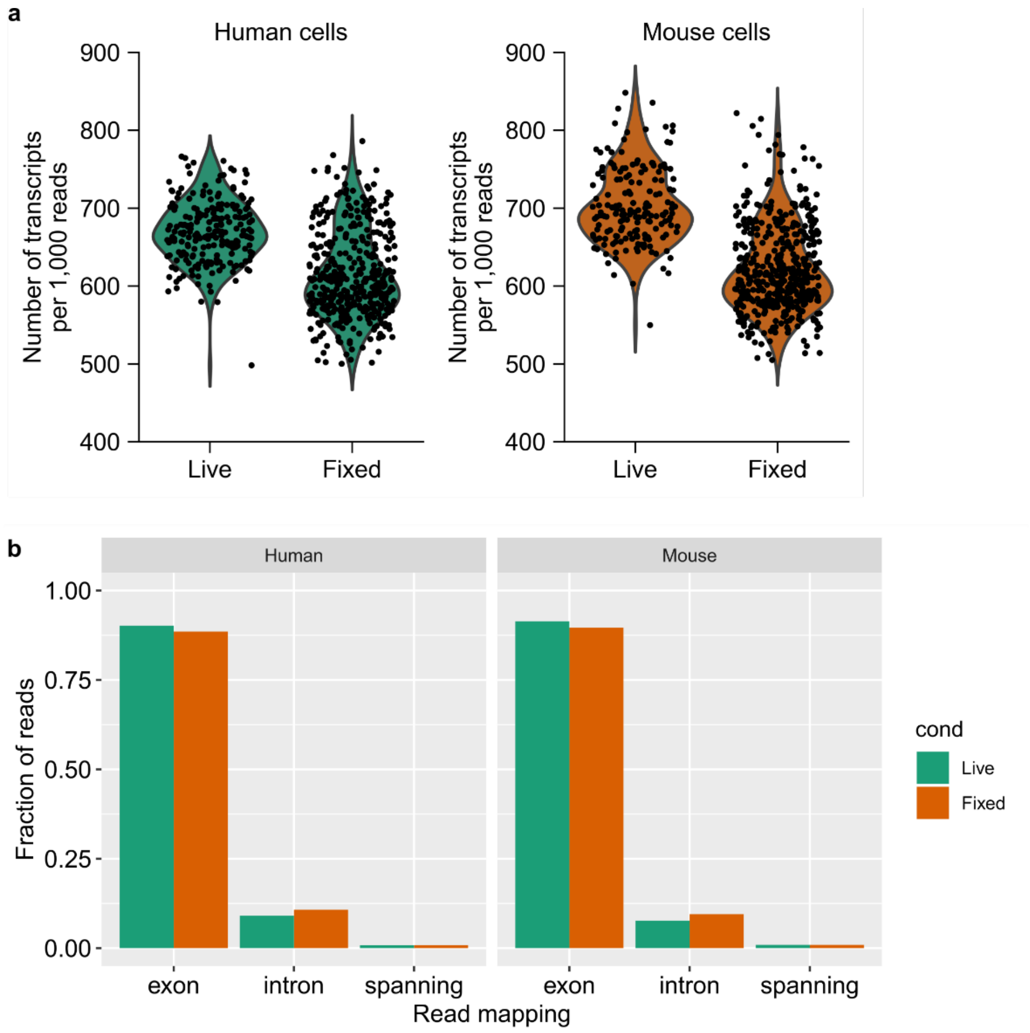


**Figure S2. Comparison between number of transcripts discovered and exon/intron mapped reads in fresh live and fixed cells. (a)** The number of transcripts per 1,000 reads detected. **(b)** The fraction of reads mapped to exon, intron, or exon/intron spanning region. The data is from the same species-mixing experiment as the main Figure 1.

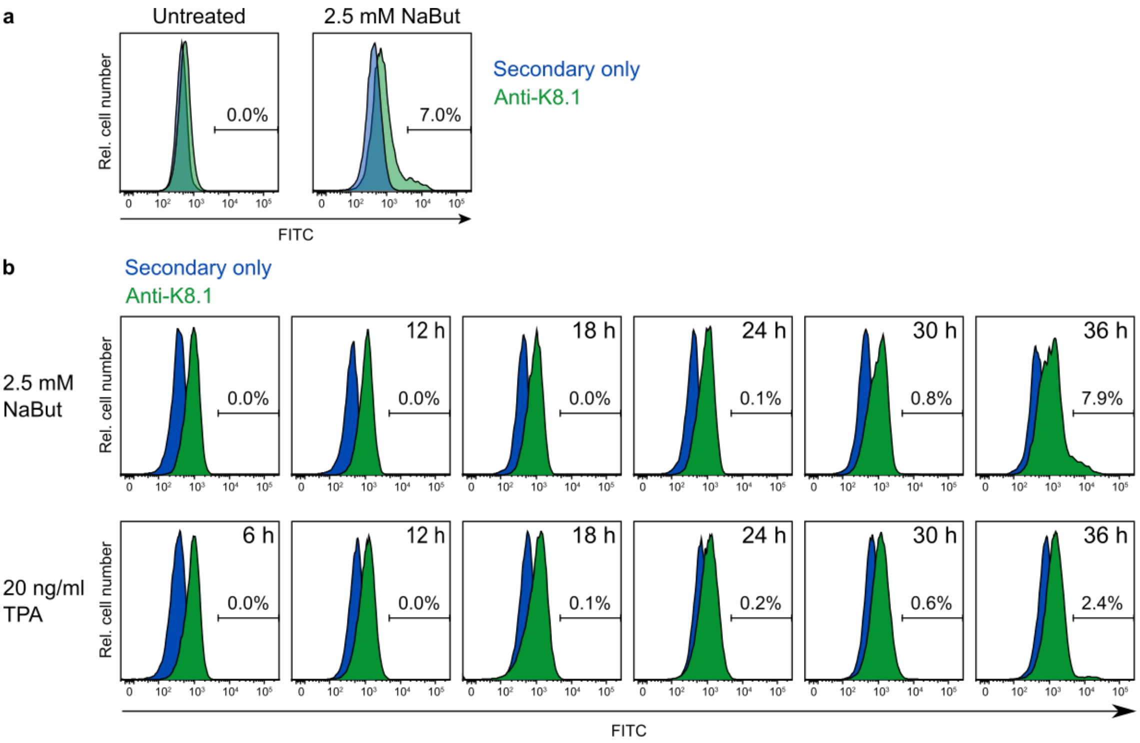


**Figure S3. Optimization of K8.1 antibody staining and induction of reactivation. (a)** Frequency of spontaneous and induced reactivation in BC3 cells. These cells were treated with 2.5 mM of NaBut for 48 hours. **(b)** Time course of reactivation induced by NaBut (top) or TPA (bottom) treatment.

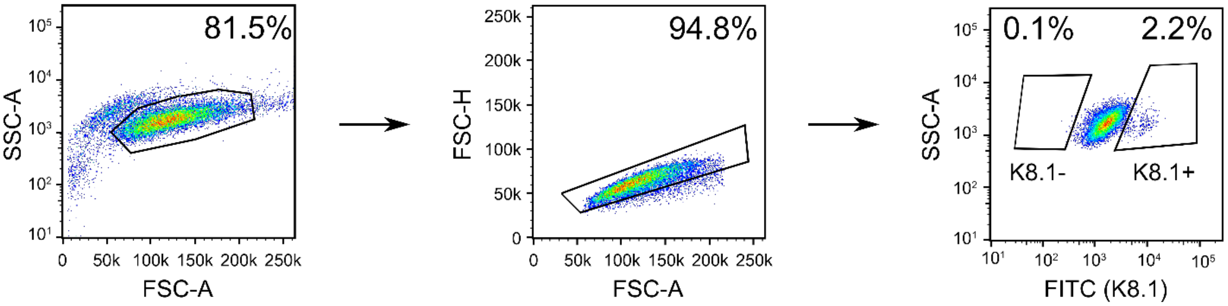


**Figure S4. Gating strategy for K8.1-positive and K8.1-negative population.**

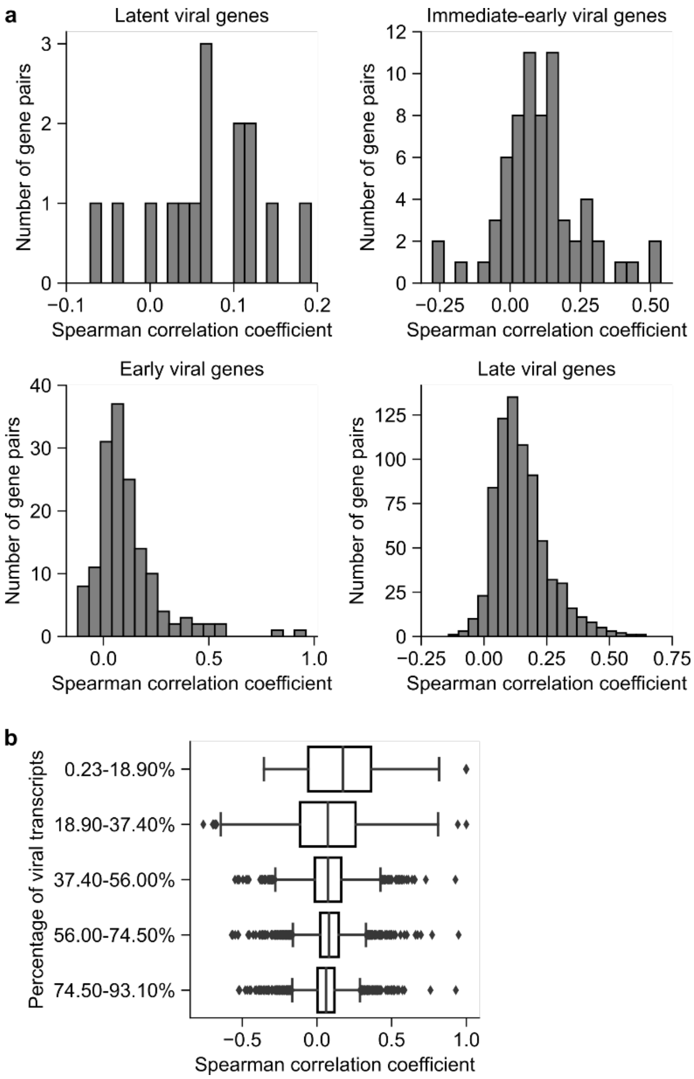


**Figure S5. Pairwise correlation of viral genes by timing and viral transcript abundance. (a)** Histograms showing the pairwise Spearman correlation coefficients of latent, immediate-early, early and late viral genes. **(b)** Box plot showing the pairwise Spearman correlation coefficient between viral transcripts binned by the percentage of viral transcripts. The middle line indicates the median of the data, the box edges indicate the first and third quartile, the whiskers indicate 1.5× the interquartile range beyond the first or third quartile, and the dots indicate the outliers.

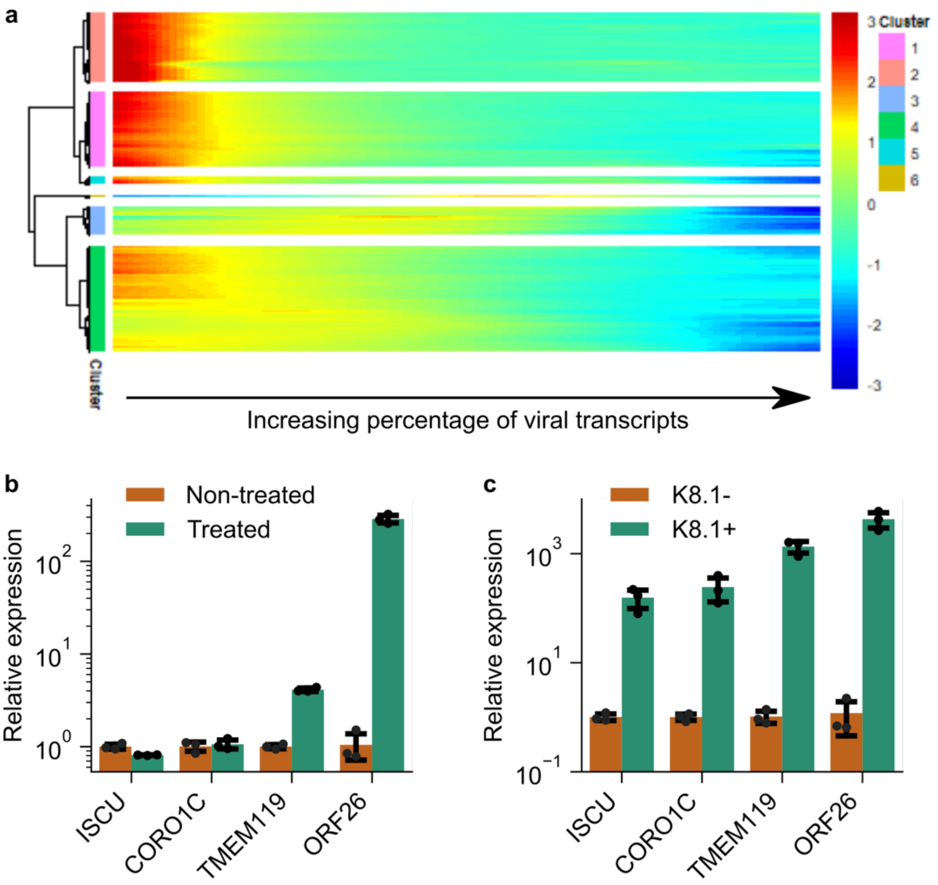


**Figure S6. Expression of host genes as a function of the abundance of viral transcripts. (a)** Heatmap of relative expression level of differentially expressed host genes. Each row shows the relative expression level of a host gene. The genes are clustered based on their expression profile, with only cluster 6 showing a positive correlation with the percentage of viral transcripts. **(b, c)** qPCR validation of the upregulated host genes in reactivated BC3 cells. The cells were treated or not treated with TPA (b), or the treated cells were sorted into K8.1-negative and K8.1 positive populations (c). (ORF26 is a viral transcript, and serves as a positive control.) n=3 biological replicates. (b,c) Data is presented as mean ± standard deviation.

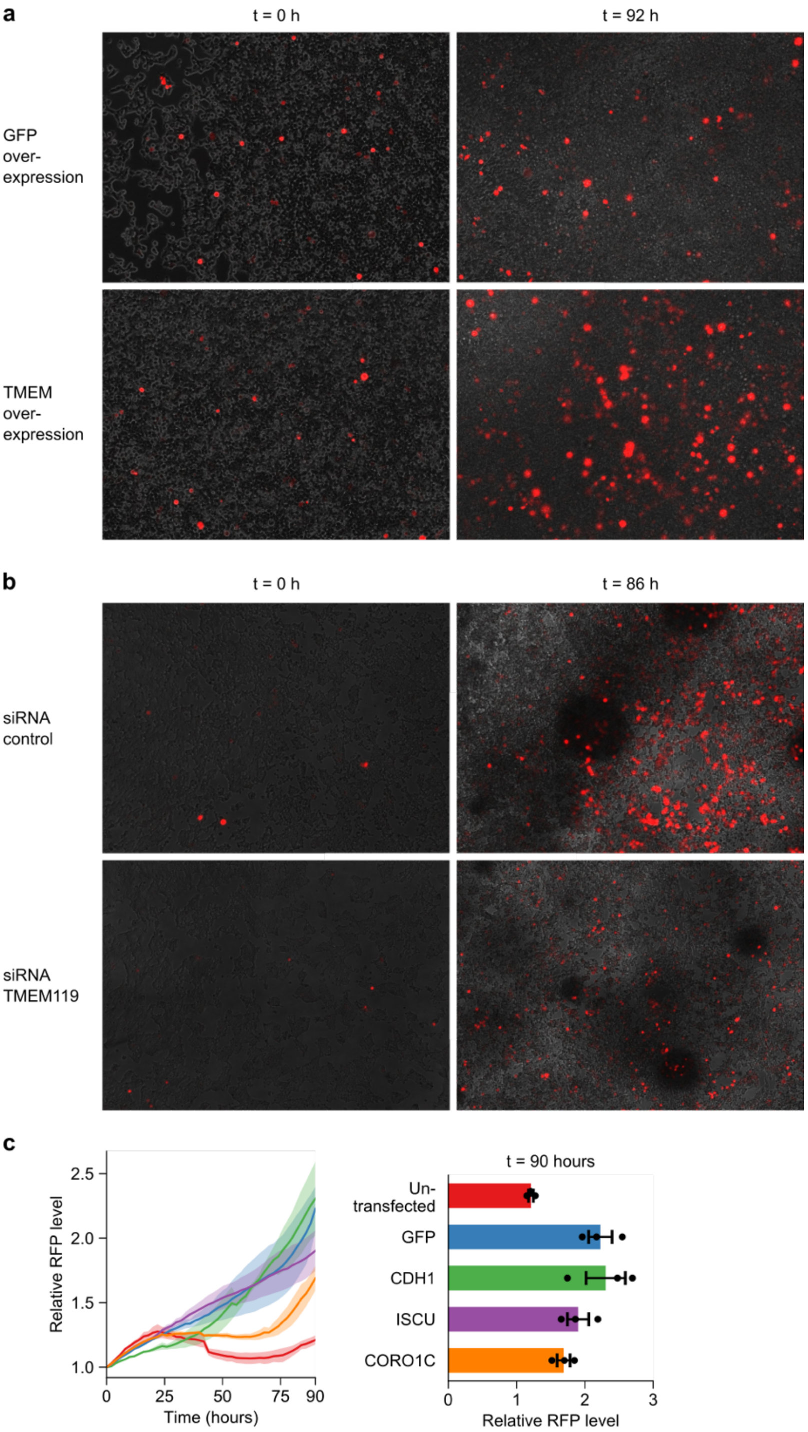


**Figure S7. Live imaging experiment. (a)** Representative images of RFP level in TPA-treated HEK293T.rKSHV219 cells transfected with GFP control (top) or *TMEM119* (bottom) at 0 and 92 hours. **(b)** Representative images of RFP level in HEK293T.rKSHV cells transfected with control siRNA (top) or TMEM119 siRNA (bottom) at 0 and 86 hours. **(c)** Time course of RFP level in HEK293T.rKSHV219 cells overexpressed with GFP control, *CDH1*, *ISCU* or *CORO1C* genes (left), and the final level at t = 90 hours (right). The data for untransfected and GFP samples are identical to the data in Figure 4b. The ribbons in the left panel and the error bars in the right panel indicate s.e.m.

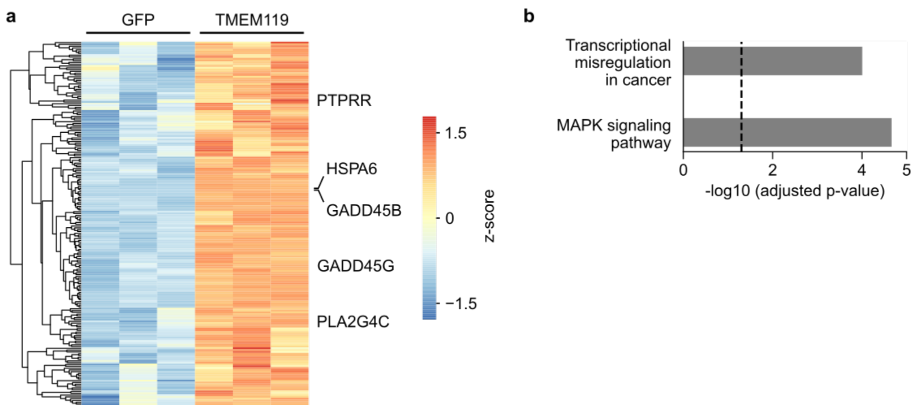


**Figure S8. Bulk RNA-seq suggests *TMEM119* modulates KSHV reactivation via MAPK signaling pathway. (a)** Heatmap showing the genes upregulated in HEK293T cells transfected with *TMEM119* compared to the GFP transfection control (n=3 biological replicates). **(b)** Bar plot showing the p-values of enriched KEGG pathways from (a). The vertical dashed line indicates P-value = 0.05.

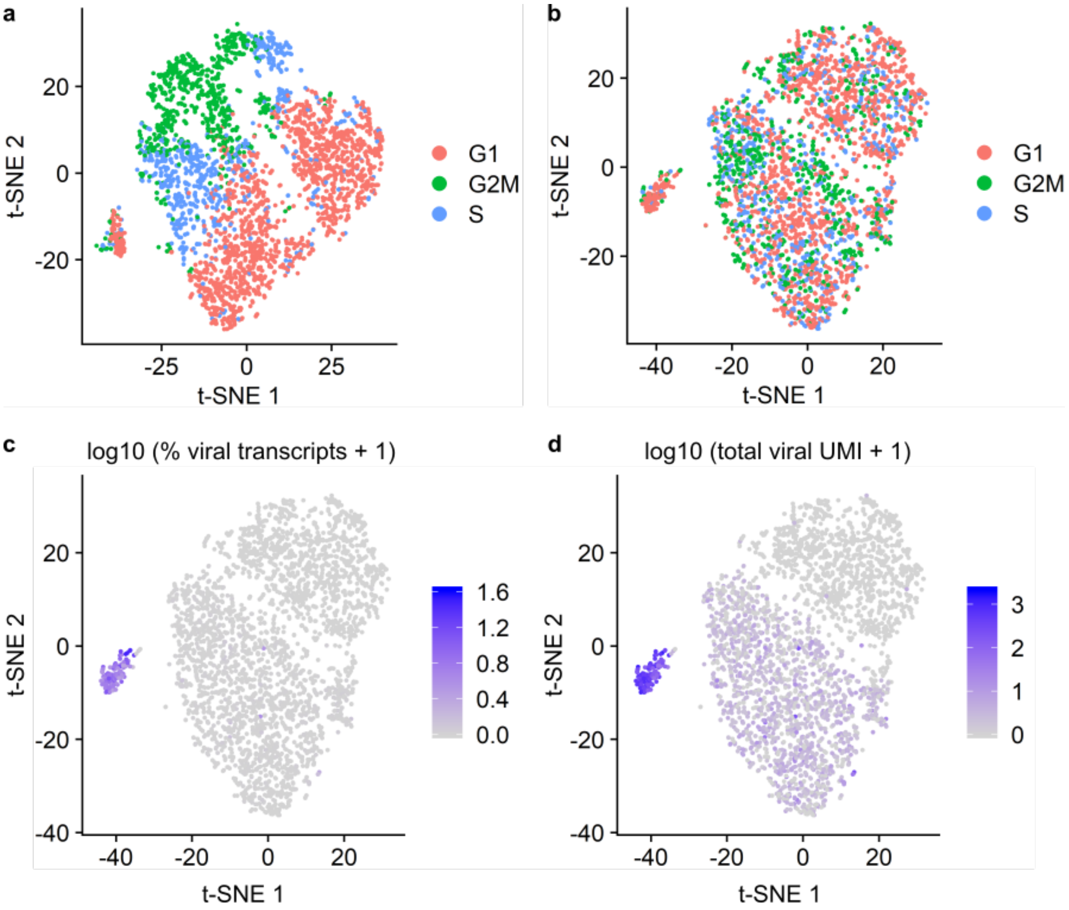


**Figure S9. Cell cycle effects and the expression of viral transcripts in OC43-infected A549 cells. (a, b)** t-SNE plots of cells (a) before and (b) after removing cell cycle effects. **(c, d)** t-SNE plots showing (c) the percentage of viral transcripts and (d) the total UMI of viral transcripts of each single cell.

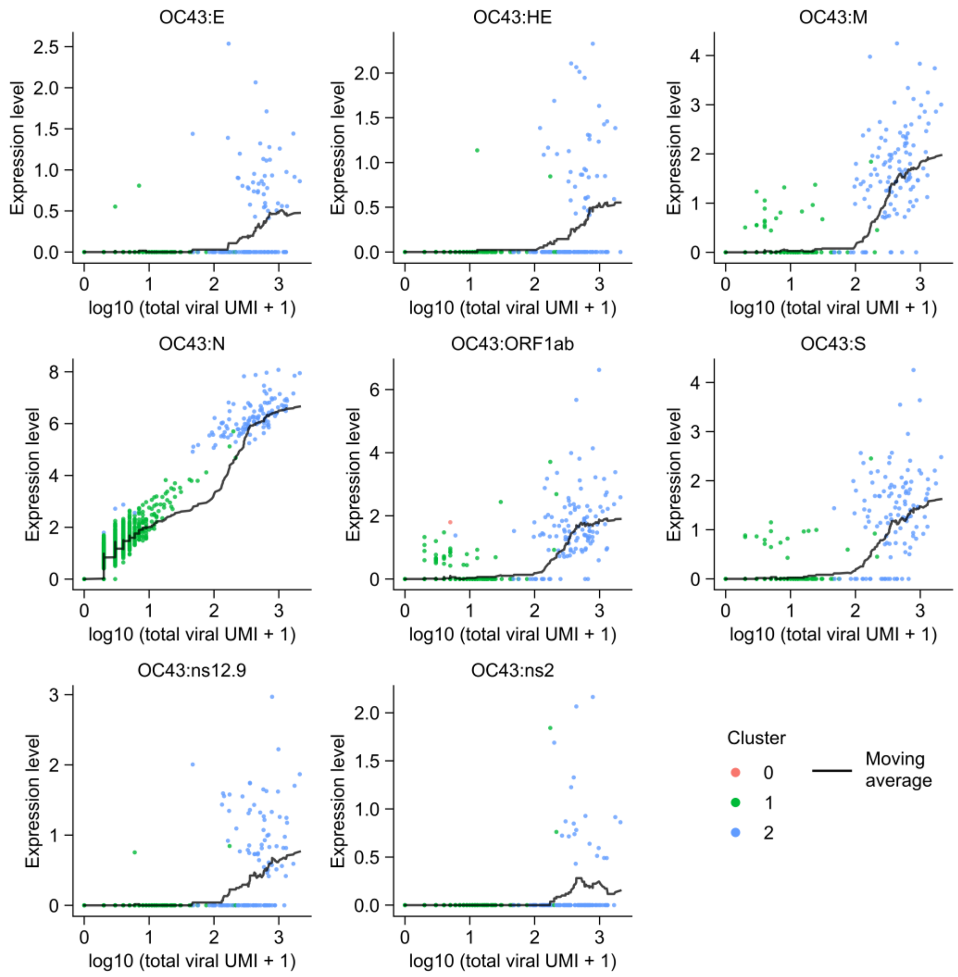


**Figure S10. Expression profiles of all OC43 viral genes in MOI 1 cells.** Scatter plots showing the expression level of each gene against the total abundance of viral genes. The black lines show the moving average of the expression level (the window is 50 cells).

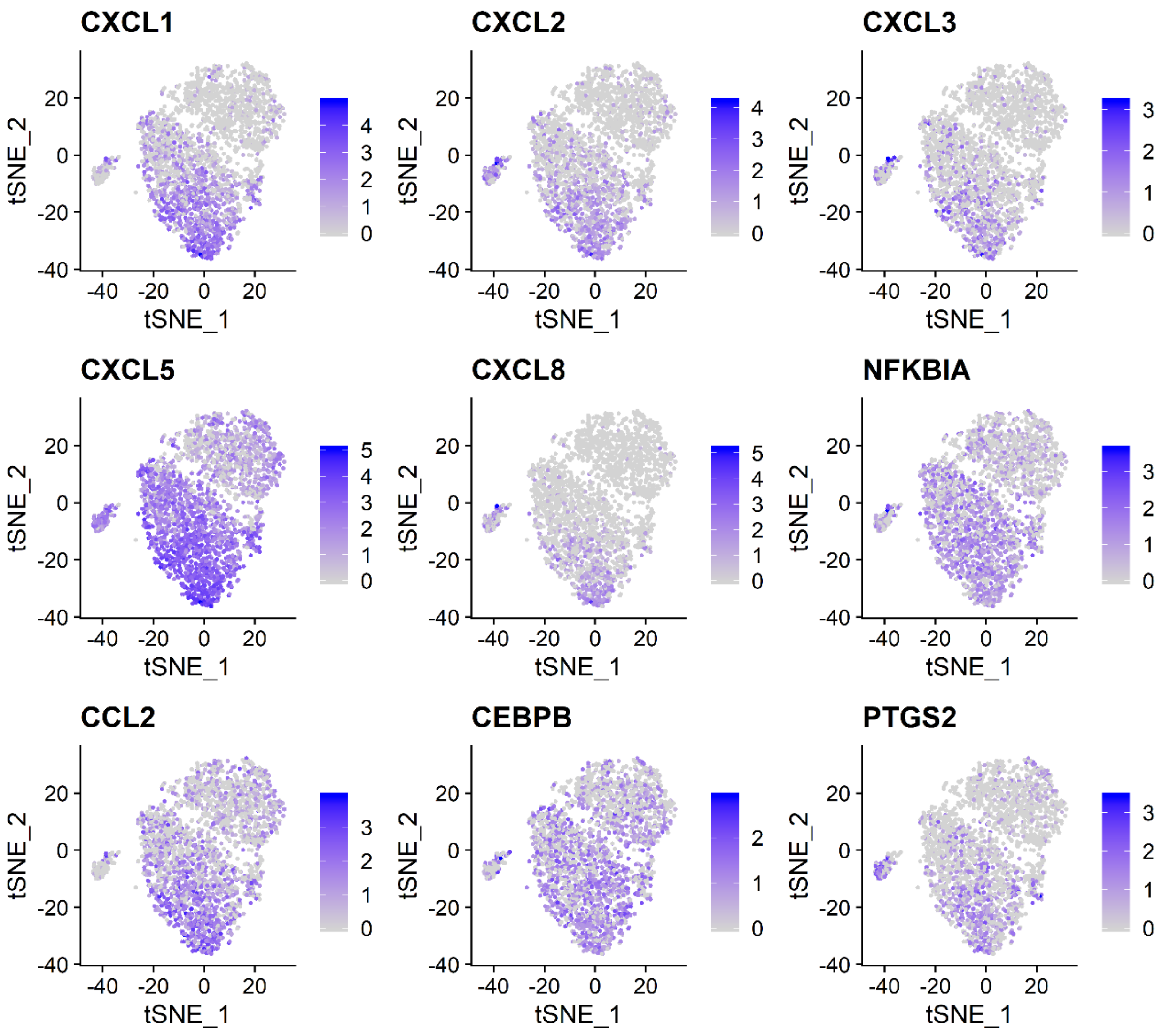


**Figure S11. Expression level of 9 representative immune-related genes.**

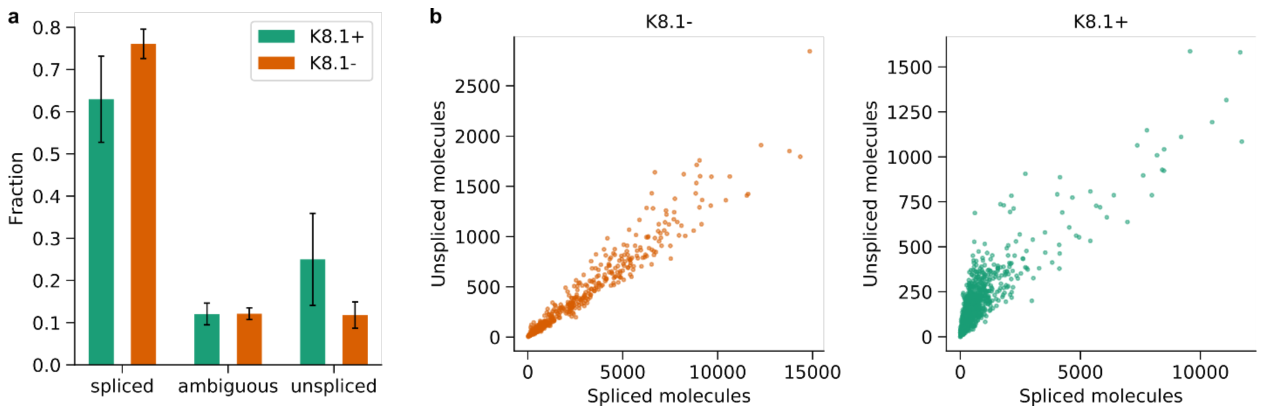


**Figure S12. Detection of unspliced host mRNAs from fixed BC3 cells. (a)** Fractions of spliced, spliced and unassigned mRNA molecules. Data is presented as mean ± standard deviation. **(b)** The number of spliced and unspliced molecules in non-reactivated (left) and reactivated cells (right).

**Table S1. Primer sequences for FD-seq**

| Name | Sequence |
|---|---|
| TSO_PCR | AAGCAGTGGTATCAACGCAGAGT |
| P5-TSO_Hybrid | AATGATACGGCGACCACCGAGATCTACACGCCTGTCCGCGGAAGCAGTGGTATCAACGCAGAGT*A*C |
| Nextera_N701 | CAAGCAGAAGACGGCATACGAGATTCGCCTTAGTCTCGTGGGCTCGG |
| Nextera_N702 | CAAGCAGAAGACGGCATACGAGATCTAGTACGGTCTCGTGGGCTCGG |
| Nextera_N703 | CAAGCAGAAGACGGCATACGAGATTTCTGCCTGTCTCGTGGGCTCGG |
| Nextera_N704 | CAAGCAGAAGACGGCATACGAGATGCTCAGGAGTCTCGTGGGCTCGG |
| Read1CustomSeqB | GCCTGTCCGCGGAAGCAGTGGTATCAACGCAGAGTAC |

**Table S2. Primer sequences for qPCR**

| Gene | Forward primer | Reverse primer | Probe |
|---|---|---|---|
| GAPDH | GAACATCATCCCTGCCTCTACTG | CAGTGAGCTTCCCGTTCAGC | |
| ISCU | CTGCACTGCTCCATGCT | CTCATTTCTTCTCTGCCTCTCC | |
| CORO1C | GTCCACTACCTCAACACATTCA | TGAAGAATCTGGCAATCTCACA | |
| TMEM119 | CCTGGCGTGAAGCAGTATTT | GCACAGGCAGAATGACACTAA | |
| ORF26 | GCTAGCAGTGCTACCCCCATT | GGTCAAATCCGTTGGATTCG | |
| ORF50 | CACAAAAATGGCGCAAGATGA | TGGTAGAGTTGGGCCTTCAGTT | AGAAGCTTCGGCGGTCCTG |
| ORF57 | TGGACATTATGAAGGGCATCCTA | CGGGTTCGGACAATTGCT | TGACGAATCGAGGGACGACGAGA |
| ORF74 (vGPCR) | GTTCCCCTGATATACTCCTGC | GGACATGAAAGACTGCCTGAG | AGGATGTACGGTCTCTTCCAAAGCC |
